# CRISPR/Cas9-based disease modelling and functional correction of Interleukin 7 Receptor alpha Severe Combined Immunodeficiency in T-lymphocytes and hematopoietic stem cells

**DOI:** 10.1101/2023.11.07.566043

**Authors:** Rajeev Rai, Zohar Steinberg, Marianna Romito, Federica Zinghirino, Yi-Ting Hu, Nathan White, Asma Naseem, Adrian J Thrasher, Giandomenico Turchiano, Alessia Cavazza

**Affiliations:** Infection, Immunity and Inflammation Teaching and Research Department, Great Ormond Street Institute of Child Health, University College London, United Kingdom; NIHR Great Ormond Street Hospital Biomedical Research Centre, London, United Kingdom

**Keywords:** Gene editing, hematopoietic stem cells, T cells, immunodeficiency, disease modelling

## Abstract

Interleukin 7 Receptor α Severe Combined Immunodeficiency (IL7R-SCID) is a life-threatening disorder caused by homozygous mutations in the *IL7RA* gene. Defective IL7R expression in humans hampers T cell precursors proliferation and differentiation during lymphopoiesis resulting in absence of T cells in newborns, who succumb to severe infections and death early after birth. Previous attempts to tackle IL7R-SCID by viral gene therapy have shown that unregulated IL7R expression predisposes to leukaemia, suggesting the application of targeted gene editing to insert a correct copy of the *IL7RA* gene in its genomic locus and mediate its physiological expression as a more feasible therapeutic approach. To this aim, we have first developed a CRISPR/Cas9-based IL7R-SCID disease modelling system that recapitulates the disease phenotype in primary human T cells and hematopoietic stem and progenitor cells (HSPCs). Then, we have designed a knock-in strategy that targets *IL7RA* exon 1 and introduces via homology directed repair a corrective, promoterless IL7RA cDNA followed by a reporter cassette through AAV6 transduction. Targeted integration of the corrective cassette in primary T cells restored IL7R expression and rescued functional downstream IL7R signalling. When applied to HSPCs further induced to differentiate into T cells in an Artificial Thymic Organoid system, our gene editing strategy overcame the T cell developmental block observed in IL7R-SCID patients, while promoting full maturation of T cells with physiological and developmentally regulated IL7R expression. Finally, genotoxicity assessment of the CRISPR/Cas9 platform in HSPCs using biased and unbiased technologies confirmed the safety of the strategy, paving the way for a new, efficient, and safe therapeutic option for IL7R-SCID patients.

## INTRODUCTION

The group of severe combined immunodeficiencies (SCID) represents the most serious form of primary immunodeficiency diseases, affecting approximately one infant in every 50,000 live births. SCID is characterized by a block in T cell development or function, variably associated with defects in B or natural killer (NK) lymphocytes. IL7R deficiency causes approximately 10% of SCID cases, and the majority of T-B+NK+ cases [1]. IL7R-SCID is caused by biallelic loss-of-function mutations in the *IL7RA* gene, which encodes for the α chain of the interleukin-7 receptor (IL7R).

The interaction of IL-7 with IL7R leads to the recruitment of intracellular signalling molecules and to activation of multiple downstream signalling pathways which are important for transcriptional activation of genes involved in T cell differentiation, survival, maturation, and TCR rearrangement. IL7R expression is restricted to T lymphocytes in humans and it is tightly regulated during T cell development to ensure correct cell maturation [2]. In the human thymus, αβ T cell thymopoiesis proceeds from haematopoietic stem and progenitor cells (HSPCs) via common lymphoid progenitors (CLPs) which commit to the T-lineage following Notch signalling. CLPs differentiate into double-negative (DN1, DN2, DN3; CD4^−^CD8^−^), then double-positive (DP; CD4^+^CD8^+^) thymocytes, which are positively selected for TCR functionality and become single-positive (SP) CD4^+^ or CD8^+^ mature T cells [3]. Only CLPs express IL7R without requiring it, allowing human *IL7RA*-deficient CLPs to develop into normal B-cells [4]. From the DN2 stage, BCL-2 expression, essential for protecting thymocytes from apoptosis, becomes IL7-dependent. IL7R also promotes V(D)J recombination in DN2–3 cells and T cell receptor γ (TCRγ) rearrangement and is thus essential for γδ T cell development [5]. IL7R expression is then downregulated at the DN4 stage and disappears in DP thymocytes, when TCR-αβ becomes expressed and takes over the anti-apoptotic role (reviewed in [2]). As such, disruption of IL-7 signalling arrests T cell development at the DN2-3 stage in *IL7RA*-knockout mice [5,6], preventing productive TCR rearrangement and leading to T cell lymphopenia. Absence of T lymphocytes results in profound failure of both humoral and cellular immunity, with severe and opportunistic infections and failure to thrive, leading to fatal outcome within the early years of life of IL7R-SCID children.

Hematopoietic stem cell transplantation (HSCT) from matched sibling donors is the leading therapy for patients with SCID, [7] however it is available to less than 20% of patients. While matched HSCT often successfully reconstitutes T cells in IL7R-SCID patients and raises their low immunoglobulin levels [8], mismatched-donor transplantation is associated with aggressive graft-versus-host disease, long-term hepatic complications of myeloablative conditioning, insufficient immune reconstitution, and mortality, especially if infection occurs [9]. Consequently, HSCT does not represent a viable therapeutic option for the remaining patients lacking a suitable donor. Genetic correction of autologous patient HSPCs and subsequent transplantation eliminates the risk of alloreactivity associated with HSCT and facilitates the use of sub-myeloablative conditioning. Successful gene therapy approaches for ADA-SCID and SCID-X1 have demonstrated the applicability of this technology to treat rare genetic diseases that affect the hematopoietic system [10]. However, pre-clinical gene therapy studies using a retroviral vector to introduce a correct copy of *IL7RA* in HSPCs derived from IL7R-deficient mice showed that constitutive, unregulated and ectopic expression of the protein can promote non-lymphoid cell proliferation, causing pre-leukemic neutrophil expansion and splenomegaly [11]. Evidence also suggests that forced IL7R expression throughout thymopoiesis diminishes the DN4 cell pool due to DP cell overconsumption and reduces thymic cellularity and peripheral T cell numbers [12]. Moreover, *IL7RA* gene amplification and human gain-of-function mutations that increase IL-7 sensitivity associate with T-ALL [13,14], while IL7R overexpression may also protect pre-leukemic T cells from apoptosis, allowing time for additional mutations to accumulate [15]. Ultimately, it has been shown that IL7R overexpression predisposes mice to thymoma [16] and inflammatory bowel disease [17]. Taken together, these observations highlight the need of tight *IL7RA* transcriptional regulation for therapeutic correction of IL7R-SCID. While the introduction of *IL7RA* specific regulatory regions in the viral vector design could serve this purpose, no studies have succeeded so far in correcting this disease through viral gene therapy, mostly due to: 1) difficulties in defining the minimum regulatory regions required to recapitulate endogenous IL7R expression; 2) the complex regulatory networks needed to control *IL7RA* transcription and translation are prohibitively large for incorporation into a viral vector to achieve tight physiological regulation; 3) semi-random vector integration pattern, which may impact on transgene expression.

An alternative to using virus-based gene therapy is to utilize genome editing, to correct the endogenous *IL7RA* locus while avoiding issues of unregulated transgene expression. Gene editing uses programmable nuclease to generate site-specific genomic double-strand breaks (DSBs) in which desired alterations can be introduced during DNA repair. The major repair pathway is non-homologous end joining (NHEJ), in which the DSB site gains a random assortment of small insertions and/or deletions (indels) and point mutations. The alternative pathway is homology-directed repair (HDR), in which a donor template with flanking arms homologous to the DSB-surrounding region accurately integrates into the DSB site. This allows targeted therapeutic transgene integration as a platform for correcting multiple mutations [18]. Robust protocols have been developed for CRISPR/Cas9 gene-editing in HSPCs [19,20,21] and several gene editing platforms to treat primary immunodeficiency diseases relying on high-fidelity HDR to integrate a therapeutic transgene in its own locus have been developed preclinically (reviewed in [22]), reaching up to 70% of HSPCs correction and complete haematopoietic reconstitution in mice, establishing the safety and efficacy of this approach.

In this study, we aim to develop a safe and effective CRISPR/Cas9-mediated genome editing platform to treat IL7R-SCID by directly knocking-in a IL7RA cDNA in frame with the endogenous *IL7RA* translational start codon, allowing regulated transcription from the promoter and enhancers naturally present in the locus, as well as the functional correction of all the mutations in the *IL7RA* gene responsible for the onset of the disease. Given the rarity of the disease and the difficulty in accessing patient blood samples, we devised a disease modelling strategy based on a multiplexed HDR platform to mimic both IL7R deficiency in T cells and HSPCs from healthy donors, as well as its restoration by monoallelic or biallelic knock-in of the corrective IL7RA cDNA through gene editing. By taking advantage of this system, we showed almost complete restoration of IL7R expression and function in IL7RA cDNA knocked-in T cells, as well as rescue of T cell development when the corrective transgene is incorporated in *IL7RA* knock-out HSPCs. Overall, this study demonstrates the feasibility and safety of a CRISPR/Cas9-based platform as a viable therapeutic approach to treat IL7R-SCID and paves the way for its potential clinical translation.

## MATERIALS AND METHODS

### Culture of human CD34+ HSPCs and T cells

Under written informed consent, CD34+ HSPCs were isolated from GCSF mobilized healthy donor apheresis using CD34+ Microbead kit (Miltenyi Biotec, UK). The percentage of purified CD34+ cell was analysed by flow cytometry staining with anti-human CD34 PE antibody (BioLegend, UK). The cells were cultured in StemSpan SFEM II medium (StemCell Technologies, UK) supplemented with Stem Cell Factor (100 ng/mL, Peprotech, UK), Thrombopoietin (50 ng/mL, Peprotech), Fms-like Tyrosin kinase 3 Ligand (100 ng/mL, Peprotech), Interleukin-3 (30 ng/mL, Peprotech), Interleukin-6 (50 ng/mL, Peprotech), StemReginin-1 (1 µM, Sigma, UK) and UM171 (35 nM, StemCell Technologies). The cells were incubated at 37°C / 5% CO_2_ for two days prior to gene editing.

To isolate T cells, human peripheral blood from healthy donors were first collected under written informed consent. After Ficoll gradient, the isolated peripheral blood mononuclear cells were cultured in X-VIVO 15 medium (Lonza, UK) supplemented with 5% human AB serum (Lonza), Interleukin-2 (50 ng/mL, Peprotech), Interleukin-7 (10 ng/mL, Peprotech) and activated by CD3-CD28 Dynabeads (ThermoFisher Scientific, UK). After incubating at 37°C/ 5% CO_2_ for three days, the Dynabeads were removed by DynaMag Magnet (ThermoFisher Scientific) before performing any gene editing experiment.

### Methylcellulose CFU assay

The colony-forming unit (CFU) assay was performed by seeding 500 cells in six-well plates containing MethoCult Enriched (StemCell Technologies) after 4 days of editing. After 14 days of incubation at 37 °C/5% CO_2_, different types of colonies including CFU-Erythroid (E), CFU-Macrophage (M), CFU-Granulocyte (G), CFU-GM and CFU-GEM were counted based on their morphological appearance.

### Selection of gRNAs

All the gRNAs used in this study, each with 20 nucleotide length sequences, were identified using the online design tool created by the Zhang lab (https://zlab.bio/guide-design-resources). Chemically modified gRNAs (Synthego, USA) contained 2-O-methyl-3′-phosphorothioate at the three terminal positions at both 5′ and 3′ ends (Synthego, USA)

### Cloning of donor templates and AAV6 production

Both KO and KI AAV6 donor vectors carrying 400 bp homology arms on each side were cloned into the pAAV-MCS plasmid containing AAV2 specific ITRs. Each of the two KO constructs comprised of either EFS-GFP (AAV6_EFS-GFP) or EFS-mCHERRY cassette (AAV6_EFS-mCHERRY) while the two IL7R corrective AAV6 contained codon optimized *IL7R* cDNA followed by either an EFS-GFP (AAV6_coIL7RA_EFS-GFP) or EFS-mCHERRY cassette (AAV6_coIL7RA_EFS-mCHERRY_KI). HEK293T cells were transfected with the AAV6 donor plasmid and pDGM6 helper plasmid, and after 72 hours the viral particles were collected by iodixanol gradient and purified using 10K MWCO Slide-A-Lyzer G2 Dialysis Cassette (ThermoFisher Scientific). AAV6 titre was determined by Quick Titre AAV quantification kit (Cell BioLabs, USA).

### Electroporation and transduction of cells

CD34+ HSPCs, human primary T cells and DG-75 cells were electroporated using the Neon Transfection kit (ThermoFisher Scientific) at 1600 volts / 10 ms / 3 pulses. The ribonucleoprotein (RNP) complex was made by combining High Fidelity Cas9 protein (IDT) with *IL7RA* targeting gRNAs at a molar ratio of 1:2 (Cas9:gRNA). To generate biallelic knockout (KO^-/-^), the electroporated cells were transduced with AAV6_EFS-GFP and AAV6_EFS-mCHERRY. For biallelic correction (KO^+/+^), the cells were transduced with AAV6_coIL7RA_EFS-GFP and AAV6_IL7R_EFS-mCHERRY whereas for monoallelic correction (KO^+/-^) cells were transduced with the AAV6_EFS-GFP and AAV6_coIL7RA_EFS-mCHERRY. The MOI used for each AAV6 was 25,000 vector genomes/cell. After 48 hours in culture, the gene edited cells were sorted by flow cytometry for the collection of highly enriched GFP^+^, mCherry+ and GFP^+^/mCherry^+^ populations.

### Artificial Thymic Organoid system

To generate Artificial Thymic Organoids (ATOs), 0.15×10^6^ of murine stromal cells expressing human delta like ligand 1 (Sigma) were combined with 7.5×10^3^ of sorted and enriched KO^-/-^, KI^+/-^ or KI^+/+^ CD34+ HSPCs. ATOs seeded with healthy donor HSPCs (WT) were used as a positive control. Each ATO pellet was resuspended in 5 µL of RB-27 medium comprising of RPMI 1640 (Thermofisher Scientific), 4% B-27 supplement (Thermofisher Scientific), 30 µM L-ascorbic acid 2-phosphate sesquimagnesium salt hydrate (Sigma), 1% penicillin-streptomycin (Thermofisher Scientific), 5 ng/mL human FLT3 ligand and 5 ng/mL human IL-7. Five µL of ATO suspension was seeded on 0.4 µm Millicell Transwell insert (Sigma) and placed in a 6-well plate containing 1 mL of RB-27 medium. The medium was changed and replaced with fresh RB-27 medium every 3-4 days. Each ATO was harvested for flow cytometry analysis at week 4, 6 and 8 post seeding.

### Flow cytometry analyses

BD FACSAria II (BD Bioscience) instrument was used for cell sorting of CD34+ HSPCs and T cells. For surface and intracellular staining, CytoFLEX (Beckman Coulter, UK) instrument was used while FlowJo v10 software (FlowJo LLC, USA) was used to analyse subsequent data. For IL7R protein detection, bulk unsorted KO^-/-^, KI^+/-^ and KI^+/+^ T cells were fixed with 1x Fixation buffer (ThermoFisher Scientific) for 30 min at room temperature. After washing, the fixed cells were permeabilised in 1x Permeabilization buffer (ThermoFisher Scientific) and stained with IL7R PE antibody (BioLegend) for 30 min at room temperature and analysed on flow cytometry. The percentage of total IL7R expression on KO^-/-^, KI^+/-^ and KI^+/+^ were determined by gating on the GFP^+^ /mCherry^+^ double positive population.

For pSTAT5 detection, bulk unsorted KO^-/-^, KI^+/-^ and KI^+/+^ T cells were cultured in X-VIVO 15 medium (Lonza) in the absence of serum and cytokines for 12 hours. After stimulation with 10 ng/mL of either IL-2 or IL-7 cytokines for 10 minutes, the cells were fixed in 4% PFA for 15 minutes at room temperature and permeabilised in Perm III buffer (BD Bioscience) for 30 minutes on ice. The cells were stained with pSTAT5 BV421 (BioLegend) antibody and analysed by flow cytometry. The precise impact of KO^-/-^, KI^+/-^ and KI^+/+^ on activated STAT5p was determined by gating on GFP^+^ /mCherry^+^ double positive population. Stimulated and unstimulated healthy donor T cells were used as positive and negative control, respectively.

### Digital droplet PCR analysis

The frequency of integrated GFP and mCherry cassettes in sorted population was quantified by Digital droplet PCR. For each reaction in a total volume of 27 µL, 60 ng of genomic DNA was combined with 1x PerfeCT Mulitplex qPCR ToughMix (Quantabio, USA), 0.05 µM Fluorescein (ThermoFisher Scientific), 1 µM each of GFP forward and reverse primer, 1 µM each of mCherry forward and reverse primers, 0.25 µM FAM labelled probe specific for GFP target amplicon, 0.25 µM HEX labelled probe specific for mCherry target amplicon, 1 µM each of reference SPDR forward and reverse primer, 0.25 µM Cy5 labelled probe specific for reference SPDR amplicon and nuclease free water. All the primers and probes were synthesized by Eurofins Genomics, Germany. Each reaction was inserted into Sapphire chips and run on Naica 3 Colour system (Stilla Technologies, France). The PCR conditions used was: 1 cycle of initial denaturation at 95°C for 10 min and 50 cycles comprising of denaturation (95°C for 30 sec), annealing (54°C for 30 sec) and extension (60°C for 6 min). After PCR, the chips were scanned and analysed by CrystalReader and CrystalMiner software (Stilla Technologies), respectively. The percentage of GFP or mCherry cassette integrated per haploid genome is calculated by the number of target amplicons positive droplets from either FAM or HEX channel divided by the number of reference amplicon positive droplets acquired from Cy5 channel.

### Validation of predicted off-target sites by deep sequencing

Potential off target sites for *IL7RA* gRNA detected by COSMID webtool [30] and GUIDE-seq were validated by high throughput next generation sequencing. CD34+ HSPCs from three healthy donors were edited with IL7R RNP complex as described above, and genomic DNA was harvested after 72h. PCR purified amplicons of 200 bp were generated from the list of off target primers. End repair, adaptor ligation and PCR indexing was performed on the denatured amplicons using NEB Next Ultra II DNA library prep kit for Illumina (New England Biolabs, UK). The resulting FASTQ files from RNP treated samples for each of the off target amplicons were analysed for indels through CRISPResso2 webtool [45] by comparing them with untreated samples.

### GUIDE-seq

Identification of potential off-target sites by GUIDE-seq [29] was performed by Creative Biogene. One million HEK293T cells were transfected with 12 µg HIFI Cas9, 4 µg *IL7RA* gRNA4 and 5 pmol of dsODN using Lonza Nucleofector 4-D (program CM-137). At 48 h post transfection, genomic DNA was extracted and sheared using a Covaris S220 Focussed-ultrasonicator to an average length of 500 bp. After end-repaired, A-tailed and ligation with adaptors containing 8-nt random molecular index, the DNA library was sequenced using Illumina Miseq. The subsequent datasets were analysed using either the *guideseq* Python package software [46].

### CAST-seq

Chromosomal aberration analysis by single targeted LM-PCR (CAST-seq) was performed and analysed as described in Turchiano et al [31]. Briefly, 1×10^6^ HSPCs were nucleofected with spCas9;gRNA4 RNP targeting *IL7RA* or mock nucleofected in duplicate. At day 4 post nucleofection genomic DNA was extracted and 500 ng of gDNA from each sample were randomly digested with NEBNext® Ultra™ II FS DNA Library Prep Kit for Illumina (NEB #E6177) to obtain fragments of ca. 350 bp and linkers were ligated. The first PCR was performed with *IL7RA* specific primer and decoy oligonucleotide together with the linker specific primer. A nested PCR was performed with the *IL7RA* and linker specific primers carrying Truseq adaptor sequences. Samples high-throughput sequencing was performed with MiSeq V2 500 cycles kit (MS-102-2003, Illumina). Data were analysed by filtering out non-specific reads utilizing the mock nucleofected sample as a control and by comparing duplicate samples. Finally, the retrieved hits were classified as chromosomal aberrations coming from the on-target specific activity (ON), spCas9 OFF-target activity (OT), homology mediated recombination events (HR) and from natural breaking sites (NBS).

### Ethics and animal approval statement

For usage of human CD34^+^ HSPC from healthy and WAS donors, informed written consent was obtained in accordance with the Declaration of Helsinki and ethical approval from the Great Ormond Street Hospital for Children NHS Foundation Trust and the Institute of Child Health Research Ethics (08/H0713/87).

## RESULTS

### Design and testing of a CRISPR/Cas9 platform to edit the *IL7RA* locus

To mediate the site-specific integration of a corrective IL7RA cDNA in the *IL7RA* genomic locus (**Figure 1A**), we designed different gRNAs targeting the first exon of the *IL7RA* gene and tested their activity in Jurkat cells. Allelic disruption (indels formation) rates of up to 84% were obtained with gRNA4 (**Supplementary Figure 1**), which was utilized for all further experiments. Delivery of the gRNA pre-complexed to a Cas9 protein as ribonucleoproteins (RNP) to peripheral blood (PB)-derived CD34+ HSPCs from healthy donors yielded up to 81% of indels formation (**Figure 1B**). To deliver the donor DNA molecule which serves as a template for HDR-mediated repair, we created an AAV6 vector that contains a GFP reporter cassette flanked by sequences homologous to the *IL7RA* genomic regions surrounding the gRNA cut site (**Figure 1C**). By RNP electroporation followed by transduction with the AAV6 donor vector, we observed targeted integration of the PGK-GFP reporter cassette in up to 52% of HSPCs, with no significant decrease in cell viability compared to mock-targeted HSPCs (**Figure 1B**-**D**). To evaluate the capacity of edited HSPCs to differentiate into multiple lineages, cells were subjected to in vitro colony forming unit (CFU) assays. Edited cells retained their clonogenic potential and produced similar frequencies of erythroid and myeloid cells without lineage skewing compared to controls, while yielding similar number of colonies of mock-treated cells (**Figure 1E**).

**Figure 1.**
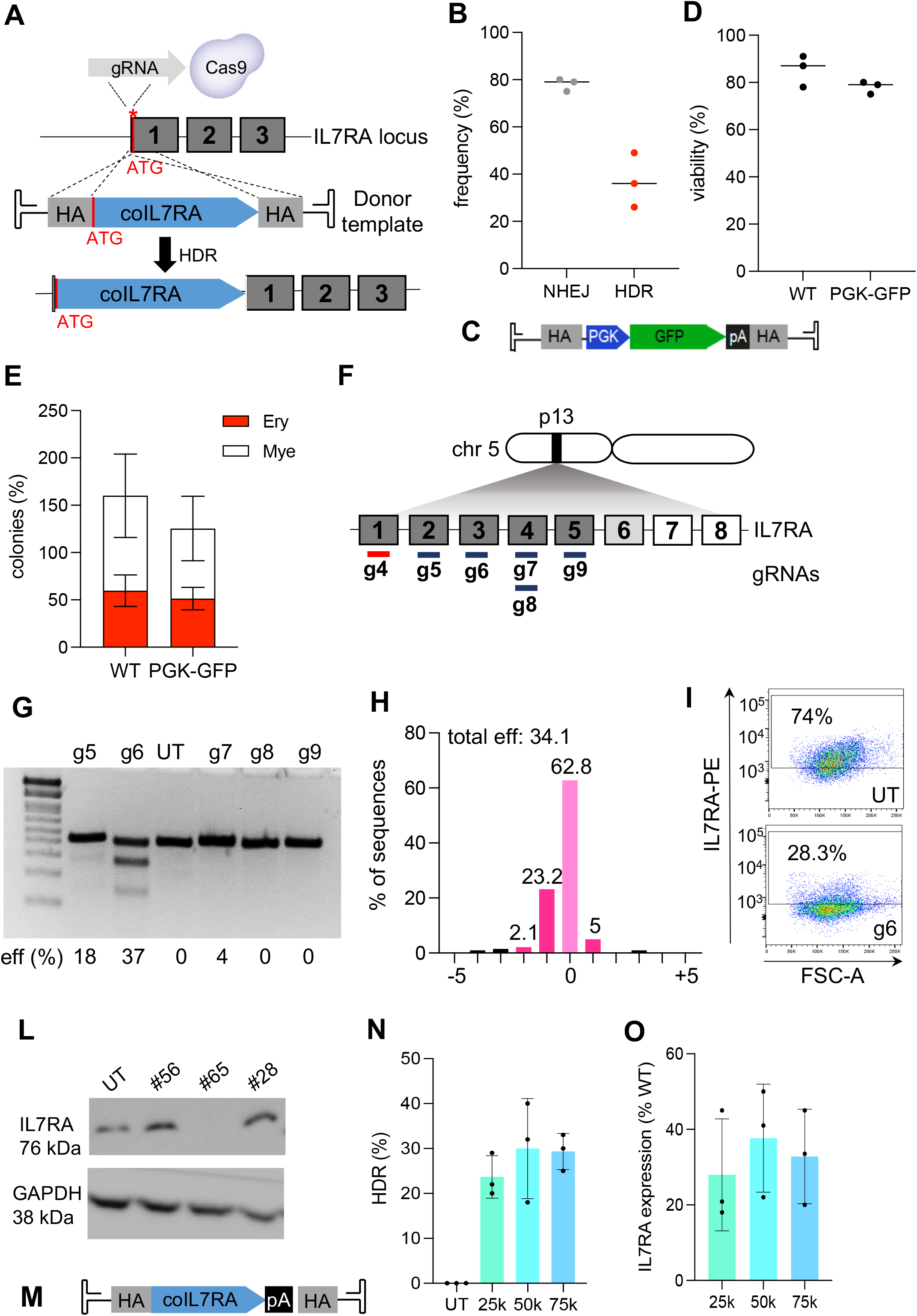
Development of a stem cell gene editing platform for IL7RA-SCID. **A)** Schematics of the gene editing strategy used to directly knock-in a codon optimized (co) IL7RA cDNA close to its endogenous promoter through HDR (HA, homology arm; ATG, translational start codon; black boxes, exons). **B and C**) HSPCs were electroporated with a Cas9:gRNA complex and indels frequency was assessed as a measure of NHEJ-mediated repair (n = 3 experiments, each dot represents a different donor source). HSPCs were also electroporated and transduced with the AAV6 donor vector containing a PGK-GFP cassette flanked by *IL7RA* HA and analysed by flow cytometry. HDR-mediated integration of the cassette can be inferred by the percentage of GFP-positive cells (n = 3 experiments using HSPCs from 3 different donor sources). **D**) Viability of non-edited (WT) and edited (PGK-GFP) cells was assessed by flow cytometry 48 hours after gene editing (n = 3 experiments using HSPCs from 3 different donor sources; p=0.25, two tailed paired Student’s *t* test). **E)** Absolute numbers of myeloid (white) and erythroid (red) colonies formed in methylcellulose by unedited (WT) and edited (PGK-GFP) HSPCs (*n* = 3 experiments from three different donor sources; p> 0.05, as analysed by one-way ANOVA with Bonferroni’s multiple comparison test). **F)** Schematics of the IL7RA gene on chromosome 5. Boxes 1-8 represents exons, where dark grey, light grey and white exons encode for the extracellular, transmembrane and intracellular IL7R domains, respectively. Lines represent the regions targeted by the gRNAs, with gRNAs 5-9 being used for gene knock out and gRNA4 being used for the gene editing strategy depicted in (**A**) to knock-in the corrective coIL7RA cDNA. **G**) Example of a T7E1 assay performed on DG-75 cells either untreated (UT) or electroporated with each of the gRNA5-9 tested (g5 to g9). The cutting efficiency (eff %) was determined by Sanger sequencing and TIDE analysis. **H**) Quantification of the size and frequency of deletions and insertions introduced by the NHEJ-mediated repair of the cut site at the *IL7RA* locus by targeted high-throughput sequencing. Fraction of sequencing reads per type of modification detected are indicated on the y axis. **I**) Quantification of IL7R expression by flow cytometry in DG-75 cells untreated (UT) or electroporated with gRNA6 (g6). **L**) IL7R protein detection by immunoblotting of DG-75 single clones sorted from DG-75 cells electroporated with the Cas9-gRNA6 RNP or untreated (UT). GAPDH expression is used as a reference. **M)** DG-75 cells were electroporated with the Cas9-gRNA4 RNP and transduced with the AAV6 donor vector carrying the corrective coIL7RA cDNA flanked by *IL7RA* HAs. **N**) Rates of targeted integration (HDR%) achieved in HSPCs after electroporation with the Cas9-gRNA4 RNP and transduced with the AAV6 donor vector at different MOIs, as assessed by ddPCR (n=3 experiments, dots represent different HSPC donor sources). **O**) Quantification of IL7R protein expression by flow cytometry in gene edited HSPCs, relative to IL7R expression detected in unedited, wild type (WT) HSPCs.

We next assessed the ability of our gene editing protocol to restore functional IL7R expression when a corrective cDNA is knocked-in at the selected site. The PGK-GFP reporter cassette in the AAV6 donor vector was therefore replaced with a codon divergent and promoterless IL7RA cDNA (coIL7RA) followed by a synthetic Bovine Growth Hormone polyadenylation (pA) signal (**Figure 1M**). Codon optimisation introduced changes in the Cas9 target sequence of the coIL7RA cDNA to prevent Cas9 from re-cutting the integrated cassette. To quantify the level of IL7R protein expressed from the coIL7RA cDNA knocked-in in frame with *IL7RA* promoter and enhancer, we developed an *in vitro* model of IL7R deficiency using a human B-lymphocyte DG-75 cell line that expresses IL7R in normal conditions. To generate *IL7RA* knock-out (KO) cell clones, we designed 5 different gRNAs targeting *IL7RA* exons 2-5 to introduce mutations that would abrogate the *IL7RA* open reading frame (**Figure 1F**). The RNP complexes containing gRNA5-9 were nucleofected in DG-75 cells and the efficacy of gene editing assessed by a T7 Endonuclease 1 (T7E1) assay and TIDE analysis, showing up to 37% of cutting frequency when gRNA6 was employed, which in turn led to a significant reduction of IL7R expression as detected by flow cytometry (**Figure 1G-1**). Clonal populations of *IL7RA* KO cells were manually selected from the edited cell bulk and assessed for IL7R deficiency by immunoblotting; clone #65 showed complete abrogation of IL7R protein expression (**Figure 1L**), further confirmed by *IL7RA* open reading frame disruption downstream the gRNA6 cut site as revealed by Sanger sequencing (**Supplementary Figure 1D**) and was thus utilized in the following experiments as an IL7R deficiency cell model. Electroporation of DG-75 clone #65 with the Cas9:gRNA4 RNP complex followed by transduction at increasing doses of the AAV6 donor template carrying the coIL7RA-pA cassette (**Figure 1M**) led to the successful insertion of the therapeutic cDNA at the *IL7RA* locus by HDR at a frequency plateauing 40% of the cells, when a multiplicity of infection (MOI) of 50,000 was used (**Figure 1N**). This corresponded to a proportional restoration of IL7R protein expression (38% of the protein level detected in wild-type DG-75 cells; **Figure 1N**), suggesting that our strategy has the potential to fully rescue physiological IL7R expression in the target cell upon knock-in of the corrective cDNA.

### Modelling and correcting IL7R-SCID in T-lymphocytes by multiplexed gene editing

Expression of the appropriate level of IL7R in target cells is critical to ensure therapeutic effectiveness. To evaluate this however, acquiring a considerable number of IL7R-SCID patient T cells or HSPCs from the peripheral blood or the bone marrow would be required, which poses challenges due to the very rare nature of the disease, the early age of IL7R-SCID infants, and the invasiveness of the procedure. To overcome this challenge, we devised an HDR multiplexing platform to mimic both *IL7RA* KO and coIL7RA cDNA monoallelic and biallelic knock-in (KI) in human primary hematopoietic cells derived from healthy donors. The *IL7RA* KO multiplexing strategy entails the use of two distinct AAV6 donor vectors carrying either an elongation factor 1α short (EFS) promoter-driven GFP or mCherry reporter cassette. The integration of these cassettes into the *IL7RA* exon 1 at the gRNA4 cut site via HDR simultaneously abrogates endogenous *IL7RA* transcription (and thus protein expression) and expresses reporter genes permitting the selection of GFP^+^/mCherry^+^ double positive cells that underwent biallelic gene KO (**Figure 2A**, **top panel**). A similar strategy allows selection of cells that bear biallelic KI of the corrective transgene, by utilising AAV6 donor vectors that carry the GFP or mCherry reporter gene preceded by the coIL7RA cDNA; the simultaneous integration of these cassettes in the *IL7RA* locus will again abrogate endogenous *IL7RA* transcription while expressing the therapeutic cDNA together with the respective reporter gene (**Figure 2A**, **bottom panel**). As monoallelic correction of the *IL7RA* locus is in principle sufficient for the disease treatment due to the functional immune systems observed in heterozygous parents of IL7R-SCID patients [8], we also paired one KO with one KI AAV6 donor vector carrying two different reporter genes, to mimic KO of the endogenous *IL7RA* gene in both alleles and KI of the corrective cassette in one allele (**Figure 2A**, **middle panel**).

**Figure 2.**
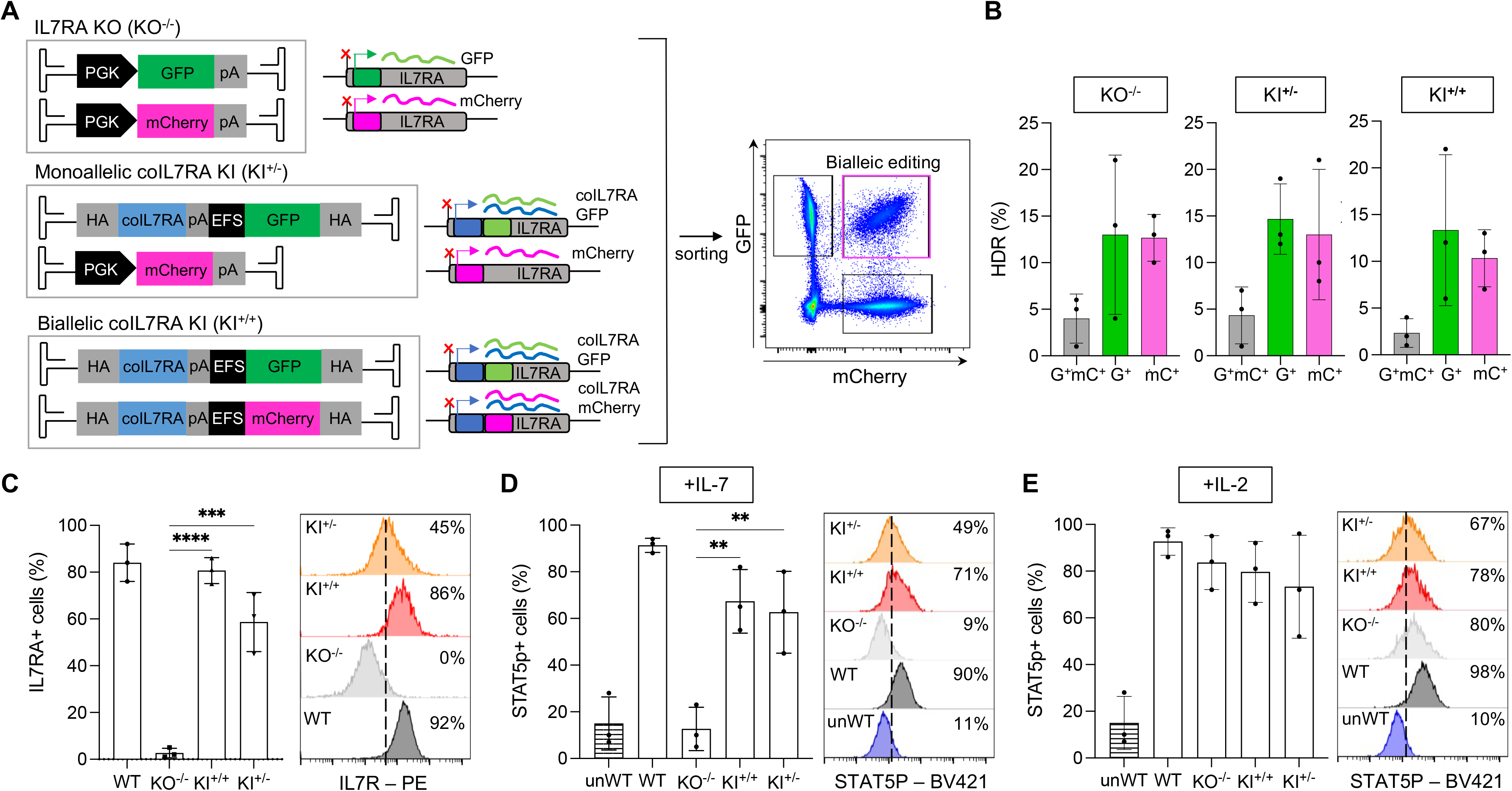
IL7R-SCID disease modelling and correction in primary T cells. **A)** Schematic of the multiplexed gene editing strategy developed to model IL7R-SCID in T cells. T cells were electroporated with a Cas9-gRNA4 RNP and transduced with a combination of two different AAV6 donor vectors to generate *IL7RA* knock-out (KO^-/-^) or coIL7RA monoallelic (KI^+/-^) or biallelic (KI^+/+^) knocked-in T cells. Pure populations of GFP/mCherry (also noted as G and mC) double or single positive cells were FACS sorted and isolated for further analysis. **B**) Frequency of targeted integration (HDR%) of the reporter cassettes in a biallelic (G+mC+) or monoallelic (either G+ or mC+ only) configuration detected in KO^-/-^, KI^+/-^ and KI^+/+^ T cells by flow cytometry (n=3 experiments, dots represent different T cell donor sources). **C**) Quantification of IL7R protein expression by flow cytometry in gene edited and unedited, wild-type (WT) T cells (left side) and representative histogram overlay plot showing IL7R expression in all groups, with the percentage of IL7R-positive cells indicated (right side) (n=3 experiments, dots represent different T cell donor sources; ****p<0.001; ***p<0.005; one-way ANOVA with Tukeys’s comparison test). **D**) Quantification of STAT5 protein phosphorylation (pSTAT5) by flow cytometry in gene edited and unedited (WT) T cells stimulated with IL-7 (left side) and representative histogram overlay plot showing pSTAT5 expression in all groups, with the percentage of pSTAT5-positive cells indicated (right side) (n=3 experiments, dots represent different T cell donor sources; **p<0.01; one-way ANOVA with Tukeys’s comparison test). **E**) Quantification of STAT5 protein phosphorylation (pSTAT5) by flow cytometry in gene edited and unedited (WT) T cells stimulated with IL-2 (left side) and representative histogram overlay plot showing pSTAT5 expression in all groups, with the fraction of pSTAT5-positive cells indicated (right side) (n=3 experiments, dots represent different T cell donor sources; p>0.5 one-way ANOVA with Tukeys’s comparison test).

T cells isolated from the PB of three different healthy donors were electroporated with the Cas9:gRNA4 RNP complex and transduced with the combination of AAV donor vectors required to achieve either complete *IL7RA* KO (KO^-/-^), monoallelic coIL7RA cDNA KI coupled with monoallelic *IL7RA* KO (KI^+/-^), or biallelic coIL7RA cDNA KI (KI^+/+^). HDR-mediated transgene integration was successfully achieved as measured by flow cytometry, with an average of 2.5-4.5% of biallelic and 10-15% of monoallelic integration per each reporter gene, respectively, totalling an average of 25-34% of editing frequency of the *IL7RA* locus across conditions (**Figure 2B**). GFP^+^/mCherry^+^ double-positive cells were FACS sorted to obtain pure populations of cells with the desired genotype; flow cytometry gating showed sufficient separation between populations for effective purification, which was further confirmed by ddPCR analysis on individually sorted populations showing correct integration of each reporter cassette in approximately 50% of the GFP^+^/mCherry^+^ double-positive cells (**Supplementary Figure 2**). To evaluate the restoration of IL7R expression in T cells in which the coIL7RA cDNA was precisely inserted in frame with the translational start site of the endogenous *IL7RA* gene, cells were fixed, permeabilised and stained for IL7R intra-and extra-cellular expression. A proportional analysis of IL7R expression compared to wildtype (WT) cells is necessary as IL7R is usually stochastically expressed in only a minority of cells even in healthy donor samples, reflecting its status as an altruistically regulated receptor [12]. Flow cytometry analysis showed a significantly increased proportion of IL7R-positive cells in WT and edited cells relative to KO^-/-^ cells, with biallelic coIL7RA cDNA KI (KI^+/+^) cells recapitulating WT expression and monoallelic KI^+/-^ cells exhibiting an intermediate proportion of IL7R-positive cells (**Figure 2C**). Comparison of IL7R mean fluorescence intensity (MFI) in edited cells showed that biallelic KI of the coIL7RA cassette led to the expression of the protein at levels comparable to those observed in WT cells, while monoallelic KI^+/-^ cells expressed it at reduced levels, likely reflecting the presence of only one functioning copy of the corrective cDNA per cell (**Figure 2C**).

To assess whether Cas9/AAV6-based integration of the coIL7RA cDNA restores IL7R functionality, we evaluated the levels of phosphorylated STAT5 protein (pSTAT5) in KI^+/-^, KI^+/+^, KO and WT T cells upon IL-7 stimulation. Indeed, binding of the IL-7 cytokine to its receptor activates a downstream IL7R signalling cascade which results in the phosphorylation of STAT5 and activation of an IL7R-dependend transcriptional profile [23]. Intracellular pSTAT5 staining in IL-7-stimulated cells indicated a trend towards restoration of the proportion of pSTAT5^+^ T cells in both biallelic and monoallelic KI cells, while KO cells showed no IL7R functionality, mimicking pSTAT5 levels detected in unstimulated IL-7 WT cells (**Figure 2D**). While STAT5 phosphorylation is a common downstream cascade effect of many cytokines mostly signalling through the common gamma chain receptor in T-lymphocytes [24], we confirmed that signalling disruption in IL7RA KO and its restoration in KI cells is strictly dependent on the presence of IL7R, as no significant difference in the frequency of pSTAT5^+^ cells was detected in all experimental groups when cells were stimulated with IL-2 (**Figure 2E**).

### Knock-in of a corrective IL7RA cDNA in HSPCs restores T cell development

Having demonstrated the successful development of a gene editing strategy to model IL7R deficiency and its correction by HDR-mediated gene knock-in in primary human T cells, we next assessed if the same approach would be effective when applied to HSPCs, the target cell type for the definitive treatment of IL7R-SCID. Because IL7R expression is neither detectable nor required in these cells, therapeutic success of the gene editing platform applied to HSPCs would be demonstrated by differentiation into mature TCRαβ^+^CD3^+^ cells, which is abrogated in the SCID phenotype. Furthermore, during the differentiation process from HSPCs to T cells, tight regulation of IL7R expression is necessary to faithfully recapitulate its physiological role during T cell development and homeostasis. To evaluate these aspects, we took advantage of the 3D Artificial Thymic Organoid (ATO) system, which models the *in vitro* differentiation and maturation of T cells from HSPCs, allowing us to elucidate the precise stages in which T cell developmental blocks occur in SCIDs and if gene editing is able to ameliorate such developmental obstructions [25,26].

To this aim, HSPCs isolated from the PB of three different healthy donors were electroporated with the Cas9:gRNA4 RNP complex and transduced with the combination of AAV donor vectors required to obtain *IL7RA* KO (KO^-/-^), monoallelic (KI^+/-^) and biallelic (KI^+/+^) coIL7RA cDNA KI. HDR-mediated transgene integration was successfully achieved as measured by flow cytometry, with a frequency of 0.4-4% of biallelic and 6-21% of monoallelic integration per each reporter gene, respectively, totalling an average of 28.4% of editing frequency of the *IL7RA* locus across conditions (**Figure 3A**). ATOs seeded with edited and FACS sorted cell populations, alongside unmanipulated wild-type HSPCs, were cultured for 6–8 weeks and their final compositions characterized by flow cytometric analysis of thymopoietic surface markers at different time points to track T cell development (**Figure 3B** and **Supplementary Figure 3A, B**). By week 4 post seeding, ATOs were able to recapitulate the two early phenotypic stages of thymic T cell progenitors: multipotent CD34+CD7−CD5− early thymic progenitors (ETP) and developmentally downstream CD34+CD7+CD5+ pro-T2 progenitors (**Figure 3C**), identified based on a classification scheme using CD5 and CD7 markers [27]. At this stage, WT, KI^+/-^, KI^+/+^ and KO^-/-^ groups show similar proportions of ETPs; however, pro-T2 progenitors are already absent from the KO^-/-^ sample, reflecting the known IL7R dependence of pro-T2 cell survival during thymopoiesis [27]. Contrary to this, the frequency of both precursor types in the monoallelic and biallelic KI samples resembled those observed in the WT samples, suggesting that knock-in of the corrective coIL7RA cDNA does rescue pro-T2 cell survival and the block in T cell development observed in patients. ATOs were further assessed for complete T cell maturation at week 6 and week 8 of culture. Similar trends as observed at earlier time points were borne out in the pre-T (CD34– CD7+ CD5+), early double positive (EDP; CD34–CD3–CD4+CD8+), and mature T cell (CD34– CD3+ TCRαβ+) populations at 6 and 8 weeks of culture, when the frequencies of these populations in the KI samples again recapitulated the behaviour of WT samples, while KO^-/-^ ones kept exhibiting curtailed thymocyte proportions (**Figure 3D** and **Supplementary Figure 3B**). These results indicate that *IL7RA* KO recapitulates the expected pro-T2 block in thymopoiesis and that coIL7R KI rescues all thymocyte populations downstream of the initial IL7R dependence at the pro-T2 stage. HAs previously discussed, IL7R expression derived from a gene addition approach must be tightly controlled throughout T cell development to avoid the occurrence of adverse events and dysregulated immunity. When looking at IL7R protein abundance in the samples seeded on ATOs, we observed correct restoration of protein expression mediated by the coIL7RA cDNA knocked-in in the *IL7RA* locus in both KI experimental groups, with biallelic KI achieving an average of 65% of WT IL7R expression in both pre-T and mature T cells. Most importantly, IL7R was not constitutively expressed throughout cell differentiation, but correctly shut down at the immature EDP stage and subsequently reactivated at the mature DP cell stage (**Figure 3E**, **F**). Overall, these data show that HDR-mediated knock-in of a *coIL7RA* cDNA in frame with its endogenous regulatory regions mediates sufficient protein expression to relieve the block in T cell development observed in IL7R-deficient patients and does so in a physiologically regulated fashion.

**Figure 3.**
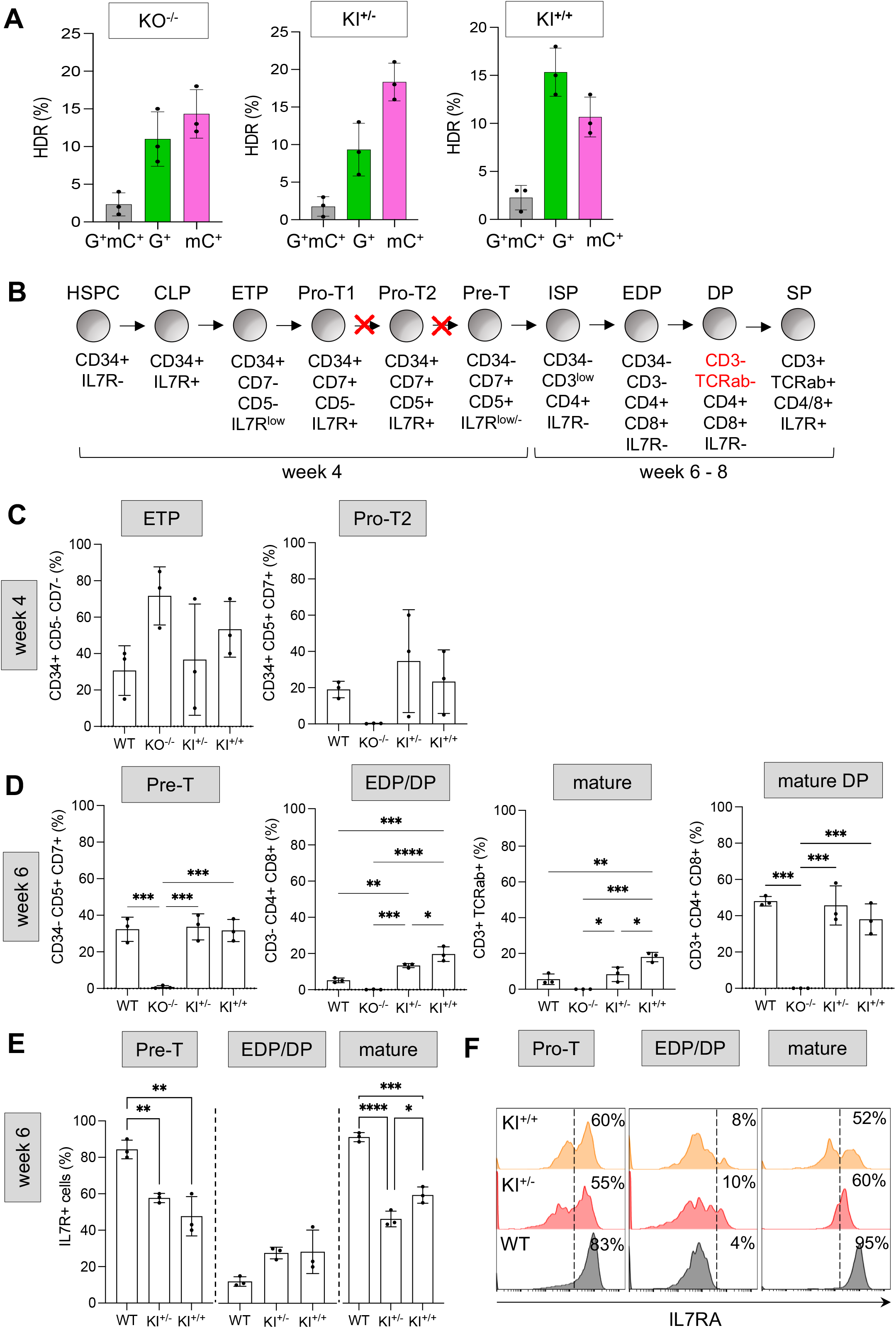
Modelling IL7R-SCID correction in HSPCs. **A)** Frequency of targeted integration (HDR%) of the reporter cassettes in a biallelic (G+mC+) or monoallelic (either G+ or mC+ only) configuration detected in KO^-/-^, KI^+/-^ and KI^+/+^ HSPCs by flow cytometry (n=3 experiments, dots represent different HSPC donor sources). **B**) Schematic of HSPC differentiation into T cells, annotated with the panel of markers relevant to detect T cell developmental subtypes in flow cytometry analysis 4-8 weeks after seeding HSPCs into artificial thymic organoids (ATOs). HSPC, haematopoietic stem and progenitor cell; CLP, common lymphoid progenitor; ETP, early thymic progenitor; Pro-T1 and Pro-T2, progenitor-T; ISP, immature single positive cell; EDP, early double-positive cell; DP, double-positive cell; SP, single-positive cell). Red crosses indicate the stages where T cell development is likely to be interrupted in IL7R-SCID patients. **C**) Frequency of ETP and Pro-T2 cells detected by flow cytometry in the different samples 4 weeks after HSPC seeding into ATOs (n=3 experiments, dots represent different HSPC donor sources; p>0.05 for all comparisons, one-way ANOVA with Tukeys’s comparison test). **D**) Frequency of Pre-T, EDP/DP, and mature DP cells detected by flow cytometry in the different samples 6 weeks after HSPC seeding into ATOs (n=3 experiments, dots represent different HSPC donor sources; ****p<0.001; ***p<0.005; ** p<0.01; *p<0.05; one-way ANOVA with Tukeys’s comparison test). **E**) Frequency of IL7R protein expressing cells in gene edited and unedited (WT) Pre-T, EDP/DP and mature T cells as detected by flow cytometry (n=3 experiments, dots represent different HSPC donor sources; ****p<0.001; ***p<0.005; ** p<0.01; *p<0.c05; one-way ANOVA with Tukeys’s comparison test). **F**) Representative histogram overlay plot showing IL7R expression in all experimental groups at the Pro-T, EDP/DP and mature T cell stages, with the percentage of IL7R-positive cells indicated on the right.

### Off-target analysis confirms the safety of the gene editing platform

One of the potential concerns associated with gene editing is the introduction of unwanted genetic modifications at off-target sites and the occurrence of gross chromosomal rearrangements, posing a huge risk for clinical therapeutic applications involving engineered nucleases [28]. To determine the specificity of our gRNA targeting *IL7RA*, we delivered the CRISPR/gRNA4 RNP to HSPCs derived from the PB of three different healthy donors and assessed the presence of indels at non-specific sites using GUIDE-seq (genome-wide, unbiased identification of DSBs enabled by sequencing) an unbiased genome-wide analysis tool [29], as well as the bioinformatic prediction tool COSMID [30]. Analysis of the GUIDE-seq sequencing data retrieved 10 different off-target sites at extremely low read numbers (**Figure 4A** and **Supplementary Table 1**). However, targeted deep sequencing of these sites in CRISPR-treated and mock-treated HSPCs from three different healthy donors demonstrated no significant gene disruption at the genomic locations tested (**Figure 4B**). In parallel, off-target activity at the top 10 genomic sites showing high homology with *IL7RA* gRNA target sequence (up to 3 mismatches) by COSMID was measured by targeted deep-sequencing in treated and mock-treated HSPCs. At a read depth of 50,000x we could not detect any genetic disruption at a statistically significant frequency compared to mock-treated controls in all genomic sites tested (**Figure 4B** and **Supplementary Table 1**). Lastly, we sought to identify the potential occurrence of large chromosomal aberrations, such as translocations, insertions or deletions, at off- and on-target gRNA cutting sites by using the CAST-seq technology [31]. For this analysis, we edited HSPCs with a Cas9:gRNA4 RNP complex containing a wild-type recombinant Cas9 protein, to be able to evaluate the presence of remote and rare rearrangement events that would not be detectable when using a high fidelity Cas9. Despite this, the CAST-seq analysis confirmed the safety of this gene editing platform, with only rearrangements at the *IL7RA* on-target locus being visible because of non-HDR mediated DNA repair at the CRISPR/Cas9 cut site (**Figure 4C**, **D**).

**Figure 4.**
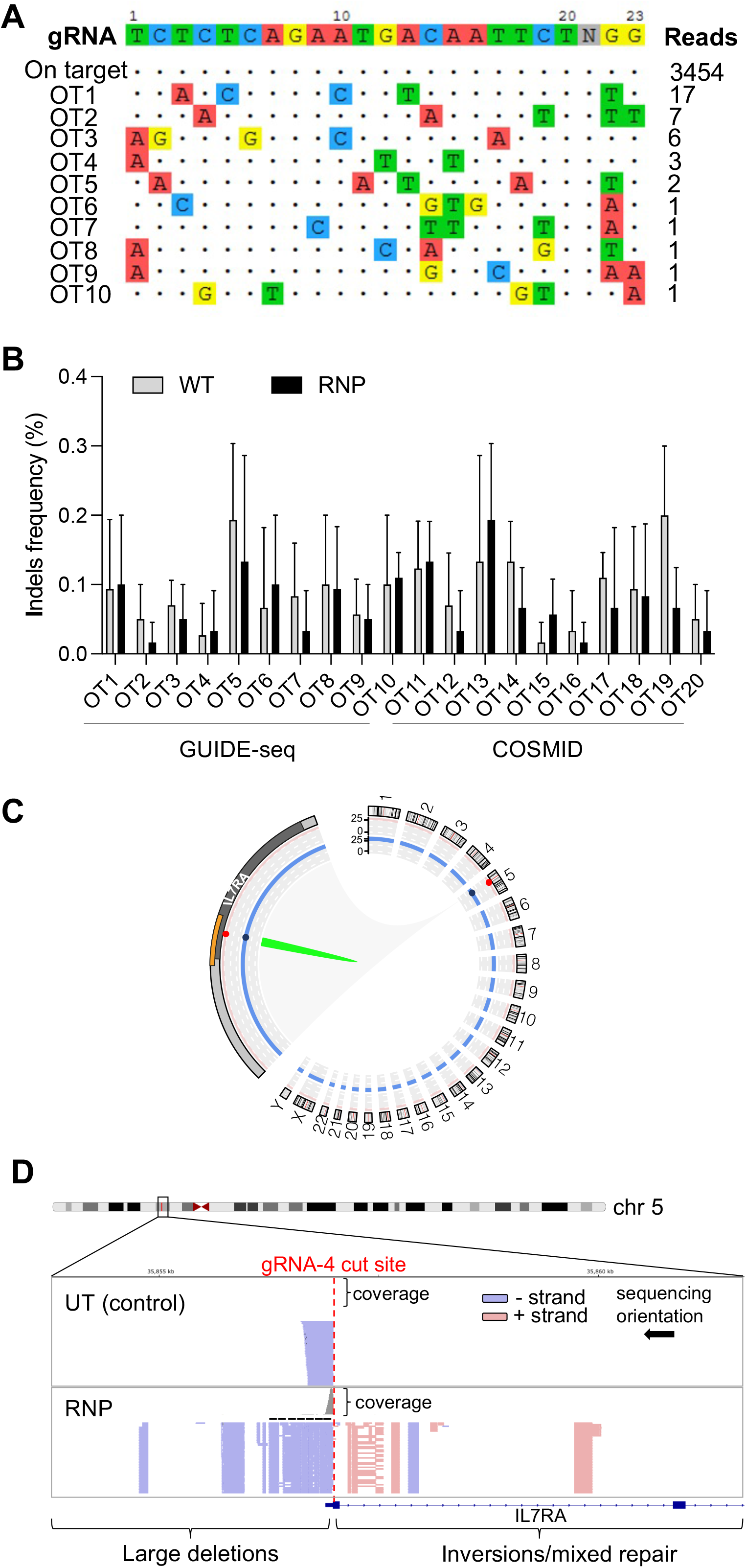
Genotoxicity analysis in edited HSPCs. **A)** Sequences of off-target sites identified by GUIDE-seq for *IL7RA* gRNA-4. The intended target sequence is shown in the top line with mismatches to the on-target site shown and highlighted in colour. GUIDE-seq sequencing read counts are shown to the right of each site. **B**) Targeted high-throughput sequencing of off-target sites detected by either GUIDE-seq or COSMID in edited (RNP) or electroporated-only (WT) HSPCs (n = 3 experiments from 3 different HSPC donor sources; no significant difference between RNP and WT samples when pooled and analysed with two-tailed paired Student’s t test). **C**) Visualization of chromosomal rearrangements detected by CAST-seq. The Circos plot shows a cluster of chromosomal rearrangement at the on-target site (green). From the outer to the inner layer, the orange rectangle shows the DNA location of the rearrangement cluster; the grey rectangle represents the *IL7RA* gene from the transcriptional start site to transcriptional termination site; a red ring indicates the alignment score against the gRNA sequence and a blue ring indicates the length of sequence homology. **D**) Qualitative CAST-Seq analysis. Integrative Genomics Viewer (IGV) plots illustrate CAST-Seq reads surrounding the target site within a window of 6 kb. Mapped CAST-Seq reads are represented by bars. Blue and red bars indicate sequences aligning with the negative or positive strand, respectively. Coverage (i.e., the number of mapped reads) is indicated in the middle and gene locations at the bottom. The positions of the on-target gRNA cutting site is emphasized by a red dotted line.

## DISCUSSION

Here, we report the successful application of a CRISPR/Cas9-based gene editing platform to model and correct a rare and devastating primary immunodeficiency, IL7R-SCID, through editing of the *IL7RA* locus in primary human T cells and HSPCs. There has been a limited number of studies showing treatment options for this disease and this work represents the first evidence of therapeutically relevant genetic correction in primary cells using a gene editing approach. IL7R-SCID represents an ideal candidate for gene editing applications for many reasons: 1) previous studies demonstrated the efficacy of genetically-corrected HSPCs in curing different forms of SCID [32,33,34]; 2) safety issues raised by preclinical studies using viral gene therapy approaches for IL7R SCID suggest the need of a physiologically regulated and restricted IL7R expression during T cell development [11,16], which can be achieved through *in situ* gene editing; 3) the tremendous selective advantage that functionally corrected cells have compared to mutated cells in a SCID setting [32,35] can compensate for the relative low HDR correction rates that can be obtained in more primitive, long-term repopulating HSCs [21] and could ideally allow infusion of autologous gene edited cells without pharmacological conditioning; 4) absence of a definitive treatment available to all the patients urges the need for new, innovative treatments to be established.

Our strategy restores IL7R expression by integrating a full-length, codon optimised IL7RA cDNA next to *IL7RA* endogenous translational start site on chromosome 5, utilising a highly specific gRNA cutting at the beginning of the *IL7RA* first exon and an AAV6 donor vector carrying a promoterless therapeutic cassette. This approach mediates the targeted integration of one/two correct copy/ies of the coIL7RA cDNA per cell, maintaining a normal gene copy number and reducing the risks of genotoxicity caused by the semi-random integration of copies in the genome of patient’s cells observed with viral gene therapy approaches [36]. Moreover, controlled integration of the corrective cDNA in the *IL7RA* locus allows the transcriptional regulation by endogenous regulatory elements, ensuring physiological *IL7RA* expression upon integration of the cassette. In addition, our platform can treat all disease-causing mutations with only one set of reagents thus representing a universal platform that could be applied to all IL7R-SCID patients and ideally to many other rare genetic disorders that require correction of the genetic defects at the HSPC level. The gene editing platform devised by us performed efficiently when tested in a *IL7RA* KO B-cell line, in T cells and in healthy human HSPCs, with high levels of targeted integration of the cassette in the desired locus and no major disruption in cell viability and HSPCs colony forming and differentiation capacity *in vitro*. However, the limited access to precious SCID patient cells limits our ability to evaluate whether expected thresholds for clinical efficacy are met in terms of editing efficiency, protein expression and function, as well as toxicity. To overcome this barrier, we have developed a multiplexing gene editing system that can efficiently mimic gene knock-out and knock-in by taking advantage of the simultaneous HDR-mediated integration of two expression cassettes with distinct reporter genes that allow FACS sorting of pure cell populations with the desired genotype. While modelling strategies similar to ours have been already devised [37,38], here we show its versatility in modelling both monoallelic and biallelic gene correction in two human primary cell types, T cells and HSPCs.

By applying our multiplexed targeted integration approach to T cells, we were able to completely abrogate IL7R expression and downstream signalling, thus obtaining a pure population of IL7R KO cells that recapitulate the IL7R-SCID defect. On the other hand, efficient monoallelic and biallelic KI of the corrective coIL7RA cDNA restored near physiological levels of protein expression, especially in the biallelic configuration, validating the use of our gene editing platform for therapeutic purposes. Assaying IL7R expression and signalling pathways is important not only for evaluating whether cDNA integration restores physiological IL7R expression and functionality, but also whether these lie below thresholds associated with oncogenic transformation risk [11,35]. Levels of phosphorylation of STAT5 are a widely used readout for IL7R signalling, as it represents a major downstream transcription factor mediating IL-7-dependent survival and differentiation programs [39]. We indeed confirmed that KI^+/+^ and KI^+/-^ cells showed significantly greater IL7R expression and pSTAT5 levels than KO T cells, while no IL7R overexpression or pSTAT5 over-phosphorylation were observed in these samples relative to WT T cells. Moreover, correction of the pSTAT5 signalling cascade by gene KI was IL7R-specific, as no changes in the signalling pathways were observed across experimental conditions when T cells were stimulated with IL-2 instead of IL-7.

Because a potential gene editing application for IL7R-SCID would require the manipulation of HSPCs as the target population, we implemented the same disease modelling strategy to those cells, in order to assess a) achievable rates of gene correction; b) functional correction of HSPCs’ capability to give rise to mature T cells once gene edited; and c) restoration of physiological protein expression in mature T cells. Evidence from bone marrow transplantation in IL7R-SCID patients and preclinical gene therapy studies have shown that as little as 10% of corrected cells engrafting the host bone marrow is sufficient to functionally cure T–B+ SCIDs due to the selective advantage of corrected cells over SCID cells [8,32].

While the efficiency of biallelic KI of the corrective cassette in HSPCs failed to meet the 10% threshold, total editing efficiencies including monoallelic KI cells frequently exceeded 25%. As such, total KI efficiency achieved is a clinically relevant figure for IL7R-SCID as, being an autosomal recessive disorder, even monoallelic correction alone is sufficient to enable normal haematopoiesis in carrier parents [8]. Previous work by our group [40] and others [20] with similar gene editing strategies applied to HSPCs reached KI rates sensibly higher than those achieved in our modelling system. We believe that this discrepancy could be due to the increased viral burden of transduction with two AAV viruses required for multiplexed HDR at high MOI, which likely artificially suppressed edited cell survival as a results of cell toxicity [41]. Moreover, the increased length of the HDR donor molecules due to the inclusion of an EFS-GFP/mCherry cassette also likely reduced KI efficiency compared to smaller clinical constructs lacking reporter genes [19,42]. This is backed up by results obtained in previous works using multiplexed HDR platforms [37,38], and additionally by the increased KI rates (up to 58%) achieved in this study when only one AAV donor vector carrying the shorter coIL7RA cDNA cassette was employed to edit HSPCs. Overall, the evidence suggests that by further counteracting gene editing- and AAV-related toxicities (e.g. by reducing the MOI used for AAV transduction, by inhibiting DNA Damage Repair pathways [42] or by optimising HSPC culture conditions [21]), we could further improve our overall KI rates *in vitro* to ensure engraftment of corrected IL7R-SCID HSPCs at therapeutically beneficial levels *in vivo*.

As the hallmark of IL7R-SCID is the immunodeficiency caused by an IL7R-dependent T-lymphocyte developmental block, we sought to understand if our gene editing platform could restore HSPCs’ ability to give rise to mature T cells *in vitro*, by taking advantage of the ATO system. The IL7RA-KO ATO model successfully recapitulated the block in thymopoiesis and absence of mature T cells seen in IL7R-SCID patients. The observation of diminished cell populations arising from differentiating KO^-/-^ HSPCs from the pro-T stage onwards agrees with previous works showing that abrogation of IL7R function causes blocks at the pro-T2/DN2 stage of thymopoiesis [6], as thymocyte survival becomes reliant on IL-7-dependent BCL-2 expression [5]. Some studies reported that this block occurs at the pre-T/DN3 stage instead, indicating there may be some variability between ATO and mouse studies, though there is broad agreement in the literature that IL7R-deficient development cannot proceed to the DP stage, as also shown by our data. On the other hand, coIL7RA-KI ATOs showed the successful overcoming of the above-mentioned developmental block when IL7R expression is reinstated, with cell frequencies detected at every developmental stage tested being comparable to WT ATOs in both mono- and bi-allelic KI conditions. Importantly, we demonstrated that IL7R expression is tightly regulated during lymphopoiesis in the ATO system, with correct down-regulation of the protein expression at the intermediate EDP/DP stages followed by its reintegration in mature TCR^+^ TCRαβ^+^ cells, highlighting the indisputable advantage of our gene editing strategy that relies on endogenous regulatory regions for gene expression. When checking the frequency of cells expressing IL7R upon gene editing and T cell differentiation, we observed an average of 65% IL7R^+^ cells (as a fraction of WT IL7R^+^ cells) in both monoallelic and biallelic KI samples at both the Pre-T and mature T cell stages analysed, exhibiting an MFI equivalent to that detected in T cells derived from WT HSPCs. The fact that we do not see an increase in IL7R expression going from monoallelic to biallelic cDNA KI suggests that there may be a maximum threshold of protein expression that can be achieved with our system, or that the populations analysed by FACS using established panels of markers contain heterogeneous population of T cells at different developmental stages and thus with different IL7R expressing levels when pushed to differentiate in the ATO system. The final, intriguing possibility is that intronless coIL7RA cDNA used in our KI strategy lacks important regulatory elements that limit transgene expression when placed under its endogenous promoter and enhancer. A previous Cas9–AAV6 gene editing approach for X-linked chronic granulomatous disease showed undetectable transgene expression using an intronless exon 1– 13 cDNA, but retention of intron 1 by downstream integration of an exon 2–13 cDNA was necessary and sufficient for physiological gene expression, presumably due to the presence of key intron-encoded regulatory regions [43]. Intronic IL7R-SCID-associated mutations have been reported as causing splicing aberrations [13], but in light of Sweeney et al.’s work they may also have unexplored regulatory significance. Future work could include bioinformatic assessment of putative regulatory regions in the IL7R introns, generation of cDNA constructs incorporating the minimal critical sequences and reassessment of KI cell function and HDR rates with the enlarged construct size. Inclusion of the ∼600-bp woodchuck hepatitis virus posttranscriptional regulatory element (WPRE) could also be an effective potential strategy for improving cDNA expression [44]. Addition of such sequences in the donor vector may however increase the size of the integrant and therefore be detrimental to the rates of targeted integration of the therapeutic cassette. Nevertheless, as heterozygous parents of SCID patients have functional immune systems, an IL7R protein expression of at least 50% of WT levels is thought be sufficient to rescue the disease phenotype, thus validating the efficacy of our gene editing platform in its current design. Moreover, the strong selective advantage of IL7R-corrected cells over deficient ones in a SCID setting may lead to eradication of the disease even when infusing HSPCs with much lower correction rates and protein expression levels than those achieved here [32]. Additionally, the lack of an increase in protein expression in biallelic versus monoallelic KI cells and relative to WT cells does imply that coIL7RA integration does not predispose T cells to overexpression of IL7R, an important safety aspects giving that IL7R overexpression associated with clonal dominance and thymocyte lymphoma [16]. While a limitation of this study is that it mostly relies on an *in vitro* assessment of the platform, it suggests that our gene-editing strategy successfully rescues thymopoiesis through IL7R expression reconstitution when applied to primary human HSPCs and has the potential to perform similarly when assessed in immunodeficient mouse models and IL7R-SCID clinical trials using patients-derived cells.

CRISPR/Cas9-mediated targeted integration through HDR provides a potentially safer strategy of gene correction than viral gene therapy, as it mitigates the risk of oncogenic transformation and genotoxicity associated with the use of viral vectors [36]. While the safety of engineered nucleases for therapeutic purposes is still under investigation, with indels generation at off-target sites being a potential threat to their safe clinical application [28], we demonstrated absence of non-specific targeting of our gRNA and of major chromosomal rearrangements upon DNA cutting, using biased and unbiased detection tools. Indeed, deep-sequencing of 20 putative off-target sites in HSPCs detected through either GUIDE-seq or a sequence similarity prediction algorithm returned no significant modifications at those sites when our optimized CRISPR/Cas9 reagents were used. This finding was further confirmed by a CAST-seq analysis in HSPCs, which showed no evident large chromosomal rearrangements between on- and putative off-target sites.

## CONCLUSIONS

Our study provides proof-of-concept of the efficacy and potential safety of a CRISPR-based gene editing approach to treat IL7R-SCID in primary human T cells and HSPCs using an *in vitro* disease modelling system. This strategy could provide a valuable therapeutic alternative for all patients affected by this disease and could enable the translation of such technology to a much wider range of HSC blood disorders.

## ACKNOWLEDGEMENTS

We thank Dr. Ayad Eddaoudi (Flow Cytometry Core Facility, University College London) for assistance with flow cytometry and cell sorting;

## AUTHOR CONTRIBUTIONS

R.R., Z.S., M.R., F.Z., Y.H., N.W., A.N., G.T. performed experiments and analysed data; A.C., G.T. and A.J.T. contributed to the study design; A.C. initiated the study, designed experiments, illustrated data, and wrote the manuscript, with inputs from all the authors.

## AVAILABILITY OF DATA AND MATERIALS

We declare that the data supporting the findings of this study are available within the paper and its Supplementary Information files or from the authors upon request.

